# Dystrophin and calcium current are decreased in cardiomyocytes expressing Cre enzyme driven by αMHC but not TNT promoter

**DOI:** 10.1101/536789

**Authors:** Ludovic Gillet, Sabrina Guichard, Maria C. Essers, Jean-Sébastien Rougier, Hugues Abriel

## Abstract

**Background:** The Cre/lox system is a potent technology to control gene expression in mouse tissues. However, cardiac alterations following cardiac-specific Cre enzyme expression in non-loxP-flanked genome heart have been reported. Recently, many loxP like sites have been identified in the wild-type mouse genome. Interestingly one of them is localized in the *Dmd* gene encoding the dystrophin protein known to be crucial for stabilization of cardiac voltage-gated ion channels Na_v_1.5 and Ca_v_1.2.

**Aim:** Here, we studied the potential alteration of dystrophin expression in adult alpha-myosin heavy chain (MHC)-Cre mice, which are extensively used for cardiac-specific recombination, and investigated Troponin T (TNT)-Cre mice as a potential alternative.

**Methods:** Cardiac-specific MHC-Cre and TNT-Cre mouse lines expressing Cre recombinase under the control of the cardiac-specific alpha-myosin-heavy chain, and rat cardiac troponin T2 promoter respectively were used. Western blots, quantitative RT-PCR, immunostainings, and patch-clamp experiments were performed to characterize MHC-Cre and TNT-Cre mouse hearts and cardiomyocytes.

**Results:** Dystrophin protein level was decreased in hearts from 12-week-old MHC-Cre^+^ mice compared to MHC-Cre^-^. Reduction of dystrophin was more pronounced with age. No significant difference was observed between 8-week-old MHC-Cre^+^ mice and MHC-Cre^-^. Immunostainings performed on cardiac sections showed reduced dystrophin signal at the lateral membrane of MHC-Cre^+^ cardiomyocytes. Quantitative RT-PCR showed decreased mRNA levels of *Dmd* gene encoding dystrophin. Finally, patch-clamp experiments showed a significant decrease in calcium current (I_CaL_) in adult MHC-Cre^+^ cardiomyocytes compared to MHC-Cre^-^. Neither dystrophin nor I_CaL_ was reduced in adult TNT-Cre^+^ mouse hearts compared to TNT-Cre^-^.

**Conclusion:** In contrary to TNT-Cre^+^ mice, the sole expression of Cre recombinase can alter the cardiac phenotype of MHC-Cre^+^ mice. Thus, researchers should include the “Cre-only” condition as control condition when designing experiments with Cre mouse strains.

## Introduction

Over the past decades, different genome manipulation technologies have been developed to investigate the function of specific genes *in vivo*. Among these, the Cre/lox system has been of particular interest^1, 2^ since it offers the possibility to generate tissue-specific and time-dependent knock-in (KI) or knock-out (KO) mice. To generate cardiomyocyte-specific KO mice, the α-myosin-heavy chain promoter, abbreviated MHC or Myh6 or αMyHC, has been largely used to control Cre recombinase expression (MHC-Cre^+^). To date, several mouse MHC-Cre^+^ strains have been generated in different genetic background. However, constitutive Cre expression in cardiomyocytes has been reported to alter cardiac functions^3^. Recently, Pugach *et al*. showed that Cre expression in MHC-Cre^+^ heart leads to a decrease of ejection fraction in an age-dependent manner and found that wild-type genome contains loxP like sites sensitive to the Cre activity^3^. Interestingly one of these loxP like site in localized in the *Dmd* gene coding for the structural protein dystrophin. Dystrophin, a 427-kilodalton protein, is a key element to maintain the integrity of the lateral membrane of cardiac myocytes^4^. The absence of dystrophin, described in dystrophinopathies, leads to alter the function of calcium and sodium voltage-gated ion channels Ca_v_1.2 and Na_v_1.5 and consequently triggers cardiac dysfunctions^5–8^. Based on these observations, we compared the dystrophin protein expression in hearts between MHC-Cre^-^ mice, MHC-Cre^+^ mice, and mice deriving from the breeding of with foxed *Dmd* gene mouse line and MHC-Cre^+^ mice. Unexpectedly, a reduction of dystrophin protein expression was observed in MHC-Cre^+^ hearts compare to MHC-Cre^-^ mice. In summary, we report on the progressive loss-of-dystrophin in one of the most used cardiac-specific Cre mouse lines, and discuss the functional consequences, in particular on cardiac ion channels.

## Methods

### Animals

All experiments involving animals were performed according to the Swiss Federal Animal Protection Law and have been approved by the Cantonal Veterinary Administration, Bern. This investigation conforms to the Guide for the Care and Use of Laboratory Animals, published by the US National Institutes of Health (NIH publication no. 85-23, revised 1996). Male mice, from three different transgenic mouse lines were used in this study. Firstly, the B6.FVB-Tg(Myh6-cre) 2182Mds/J, also known as MHC-Cre, in which the cardiac-specific α-myosin-heavy chain (Myh6) promoter drives expression of Cre was purchased from The Jackson Laboratory (Stock No 011038). Secondly, the STOCK Tg (Tnnt2-cre) 5Blh/JiaoJ also called cTnT-Cre where Cre recombinase expression is under the control of the rat cardiac troponin T2 promoter was purchased from the Jackson Laboratory (Stock No 024240). From a mixed background, these mice were bred to C57BL/6J for at least one generation upon arrival at the Jackson Laboratory. Thirdly, the *Dmd*^flox^ mice in which exon 22 is flanked by loxP sites was generated by PolyGene AG (Rümlang, Switzerland).

### Protein extraction and western blots

Whole hearts were extracted from mice, the atria were removed and the ventricles were homogenized in 1 ml lysis buffer (50 mM HEPES, 150 mM NaCl, 1 mM EGTA, 10% glycerol, 1.5 mM MgCl2, and Complete^®^ protease inhibitor cocktail (Roche, Basel, Switzerland) using a Polytron. Triton X-100 in lysis buffer was then added to make a final concentration of 1% (final lysate volume 2 ml). Samples were lysed on a rotating wheel for 1 hour at 4°C. Soluble fractions of the mouse heart were obtained by subsequent centrifugation at 13,000 rpm for 15 minutes at 4°C. To be able to load esimilar protein amounts on a gel, protein concentrations were measured in triplicate by Bradford assay and extrapolated by a bovine serum albumin (BSA) standard curve. Samples were denatured at 95°C for 5 minutes prior to gel loading. To analyse and compare protein expression, 80 μg of each ventricular mouse heart lysate sample were loaded on a 1.5 mm-thick homemade SDS Page gel either 10% or 8-15% for high molecular weight proteins. Gels were run at 150 V for 1 hour and the protein were subsequently transferred to nitrocellulose membranes using the Biorad Turbo Transfer System (Biorad, Hercules, CA, USA). All membranes were stained with Ponceau as a qualitative check for equivalent of total protein loading and transfer. Membranes were then rinsed twice with 1X PBS before using the SNAP i.d. system (Millipore, Zug, Switzerland) for western blotting. One % Bovine Serum Albumin (BSA; Sigma) was used as blocking solution and incubated with membranes for 10 minutes before incubation with primary antibody for 10 minutes. Membranes were subsequently washed 4 times in 1X PBS + 0.1% Tween before incubating with secondary fluorescence antibodies for 10 minutes. After 4 more washings with 1X PBS + 0.1% Tween and 3 washings in 1X PBS, membranes were scanned with the Odyssey^®^ Infrared Imaging System (LI-COR Biosciences, Bad Homberg, Germany) for detection of fluorescent protein. Subsequent quantitative analysis of protein content was achieved by measurement and comparison of band densities (equivalent to fluorescence intensities of the bands) using the Odyssey Version 3.0.21 software.

### Immunohistochemistry of mouse ventricular sections

Following removal of atria, hearts were cut transversally, embedded in Tissue-Tek^®^ O.C.T™ Compound (Sakura Finetek, Zoeterwoude, the Netherlands), and cryo-preserved. Frozen sections, 10μm-thick, were stored at −80°C until later use. Sections were then thawed at room temperature for 30 minutes and subsequently fixed in cold acetone for 10 minutes. Samples were dried at room temperature for 10 minutes. After rinsing in 1X PBS, sections were blocked in 1X PBS with 0.5% Triton, 1% BSA, and 10% goat serum for 30 minutes. Sections were rinsed again in 1X PBS before addition of primary antibodies (in 1X PBS with 0.5% Triton, 1% BSA, and 3% goat serum). After 2 hours of incubation, sections were rinsed again in 1X PBS and then incubated for 1 hour in secondary antibody. After a final wash with 1X PBS, FluorSave™ Reagent (Calbiochem, La Jolla, CA, USA) was applied to the sections, which were then stored at 4°C and analysis using confocal microscope (Zeiss Z-710).

### Antibodies

For western blots, a mouse monoclonal against dystrophin (Dys NCL-DYS2, Novocastra, Muttenz, Switzerland) was used at the dilution of 1/250. A custom-made rabbit polyclonal antibody against syntrophin (Pineda Antibody. Service, Berlin) was used at a dilution of 1/1000. A mouse monoclonal antibody against SAP97 (VAM-PS005F, Enzo life science, Lausen, Switzerland) was used at a dilution of 1/500. Against Na_v_1.5, a custom rabbit polyclonal antibody (Pineda Antibody. Service, Berlin) was used at a dilution of 1/200. A rabbit polyclonal antibody against Cre (69050; Novagen, EMD Millipore-MERK, Schaffhausen, Switzerland) was used at a dilution of 1/500 and rabbit polyclonal antibody against calnexin (C4731; Sigma-Aldrich, Saint-Louis Missouri, USA) was used at 1/1000.

For immunostainings rabbit polyclonal antibody against Cre (69050; Novagen, EMD Millipore-MERK, Schaffhausen, Switzerland) was used at a dilution of 1/500 and mouse monoclonal anti-dystrophin (Dys NCL-DYS2, Novocastra, Muttenz, Switzerland) was used at a dilution of 1/250.

### RNA extraction and quantitative RT-PCR

Total RNA was isolated from mouse heart pieces, with the ExpressArt RNAready kit (Amp Tec GmbH, Germany, provided by the lab of Pr. R. Jaggi, Bern, Switzerland). The High Capacity cDNA Reverse Transcription Kit (Applied Biosystems, Life Technologies, Switzerland) was used to synthesize cDNA. Subsequently, 25 ng of cDNA was mixed with 1X TaqMan Universal Master Mix (Invitrogen) and 1 µL of either dystrophin (*Dmd*) or glyceraldehyde-3-phosphate dehydrogenase (GAPDH) probe/primer mix (Applied Biosystems; Mm00464475_m1 and Mm99999915_g1, respectively). GAPDH was used as the reference gene to which experimental data was normalized. All samples were loaded in quadruplicates. On an ABI 7500 RT-PCR machine, samples were run with the following thermal cycling conditions: holding stage at 95°C for 20 seconds, then 40 cycles of 95°C for 3 seconds and 60°C for 30 seconds. Relative *Dmd* expression (ΔCT) was calculated by subtracting the signal threshold cycle (CT) of the control (GAPDH) from the CT value of Dmd. Subsequently, ΔΔCT values were obtained for MHC-Cre mice by subtracting the ΔCT of the control mice from the ΔCT of MHC-Cre mice. Results were then linearized by calculating 2^-expΔΔCT^.

### Isolation of mouse ventricular myocytes

Single cardiomyocytes were isolated according to a modified procedure of established enzymatic methods. Briefly, mice were euthanized by cervical dislocation. Hearts were rapidly excised, cannulated and mounted on a Langendorff column for retrograde perfusion at 37°C. Hearts were rinsed free of blood with a nominally Ca^2+^-free solution containing (in mM): 135 NaCl, 4 KCl, 1.2 MgCl_2_, 1.2 NaH_2_ PO_4_, 10 HEPES, 11 glucose, pH 7.4 (NaOH adjusted), and subsequently digested by a solution supplemented with 50 µM Ca^2+^ and collagenase type II (1 mg/mL, Worthington, Allschwil, Switzerland) for 15 minutes. Following digestion, the atria were removed and the ventricles transferred to nominally Ca^2+^-free solution, where they were minced into small pieces. Single cardiac myocytes were liberated by gentle trituration of the digested ventricular tissue and filtered through a 100 µm nylon mesh. Ventricular mouse cardiomyocytes were used after an extracellular calcium increase procedure to avoid calcium overload when applying extracellular solutions in electrophysiology assays.

### Electrophysiology

Whole-cell currents were measured at room temperature (22–23°C) using a VE-2 amplifier (Alembic Instrument, USA). The internal pipette solution was composed of (in mM) 60 CsCl, 70 Cs-aspartate, 1 MgCl_2_, 10 HEPES, 11 EGTA and 5 Mg-ATP, pH 7.2, with CsOH. The external solution contained (in mM) 130 NMDG-Cl, 5 CsCl, 2 CaCl_2_, 1.2 MgCl_2_, 10 HEPES and 5 D-glucose, pH 7.4, with CsOH. Data were analysed using pClamp software, version 10.2 (Axon Instruments, Union City, California, USA). Calcium current densities (pA/pF) were calculated dividing the peak current by the cell capacitance. Activation curves and steadystate inactivation curves were fitted with the following single Boltzmann equation: y=1/ (1+exp [(*V*h–*V*_50_)/k]), in which *y* is the normalized conductance or peak current at a given holding potential (*V*h); *V*_50_ is the voltage at which half of the channels are activated (*V*_50_, act) or inactivated (*V*_50_, inact) respectively, and *k* is the slope factor.

### Statistical Analysis

Data are represented as means ± S.E.M. Statistical analyses were performed using Prism7 GraphPad™ software. Unless otherwise specified in the figure legends, a Mann-Whitney two-tailed U test was used to compare two groups. *p*<0.05 was considered significant. N referees to number of mice/hearts used and n to number of isolated cardiomyocytes used.

## Results

### 3.1 Dystrophin protein expression is reduced in age-dependently manner in MHC-Cre mouse hearts

As briefly described in the methods section, *Dmd*^flox+^ mice have been generated and used as a control for the Cre recombinase activity. Western blots were performed using heart lysates from wild-type (WT) (*Dmd*^flox-^ / MHC-Cre^-^), MHC-Cre^+^ (*Dmd*^flox-^/MHC-Cre^+^), and dystrophin KO (*Dmd*^flox+^ /MHC-Cre^+^) littermate mice. As shown in figure 1A, dystrophin expression was efficiently suppressed in dystrophin KO compared to WT hearts. Surprisingly, in the non-floxed *Dmd* gene MHC-Cre^+^ littermates, also used as controls, dystrophin expression was greatly reduced (Fig. 1A, compare lane 2 and 6 with lane 3-5). In contrary, the presence of the floxed element in *Dmd* gene only did not affect the dystrophin expression (Suppl. Fig. 1A). To address the question of the time course of this observation, western blots were performed on hearts from 8-, 10-, 12-, and 26-week-old mice. As shown in figure 1B, the decrease of dystrophin protein was more pronounced with age. To confirm that the loss of dystrophin expression was age-dependent, western blots were performed on 8- and 12-week-old mouse hearts from MHC-Cre^+^ and control littermates (MHC-Cre^-^). While dystrophin expression did not differ between control and MHC-Cre^+^ hearts at 8 weeks, at 12 weeks dystrophin expression was significantly decreased by 50 ± 10% in MHC-Cre^+^ hearts compared to the respective controls (Fig. 2A and 2B). Finally, immunostainings on cardiac sections of 12-week-old mouse hearts confirmed the decrease of dystrophin expression at the lateral membrane of MHC-Cre^+^ isolated ventricular cardiomyocytes compared to their littermate controls (MHC-Cre^-^) (Fig. 3A).

**Figure 1.**
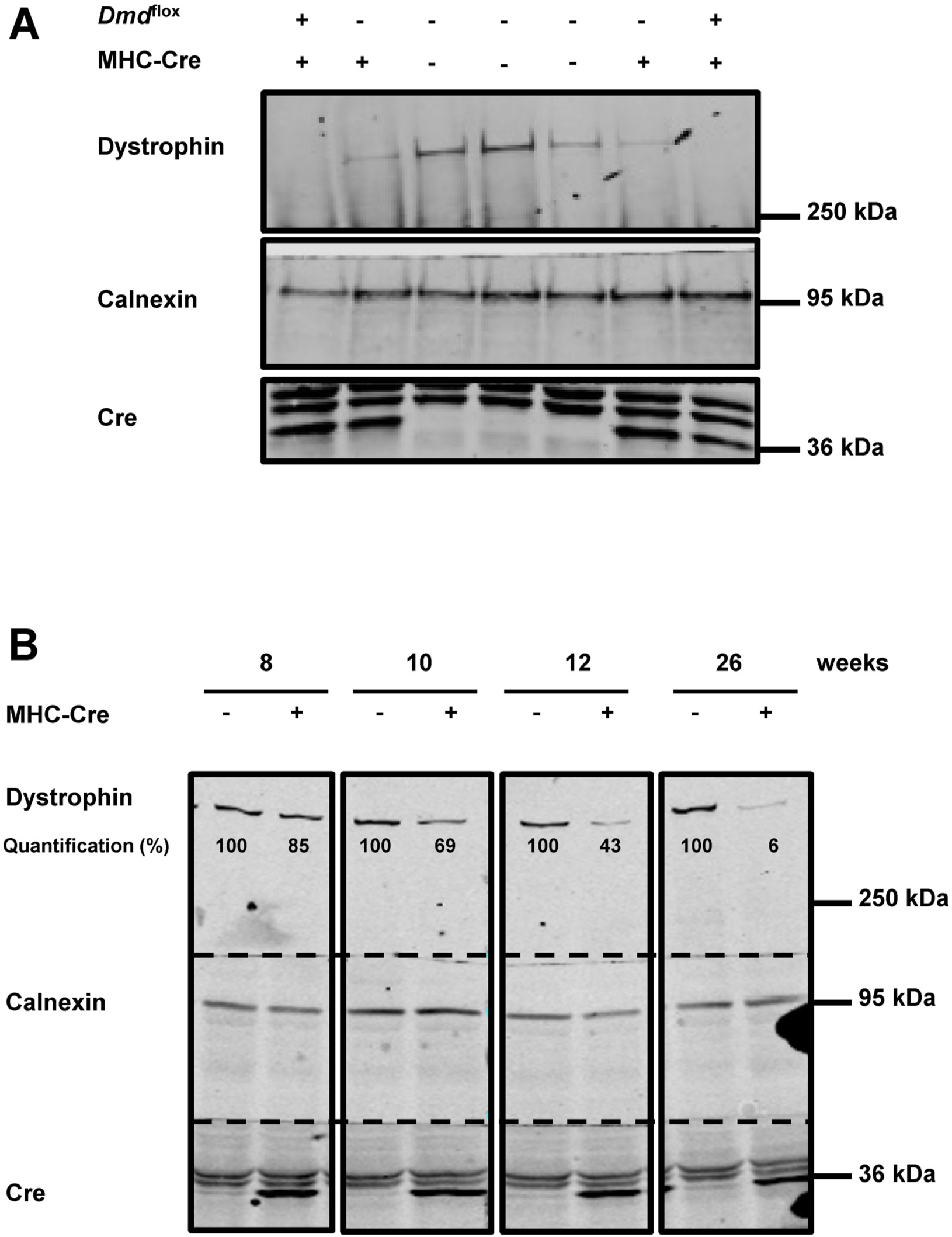
Dystrophin expression is decreased in adult mouse hearts expressing Cre. A) Western blots showing dystrophin expression in the heart of WT (*Dmd*^fIox-^, MHC-Cre^-^), MHC-Cre^+^, and cardiac-specific dystrophin-KO (*Dmd*^fIox+^, MHC-Cre^+^ mice) (N=2 to 3 for each condition). B) Western blots showing dystrophin expression in mouse hearts at different ages with or without Cre recombinase expression. Dystrophin expression is quantified by normalizing band intensity to calnexin expression (N=1 for each condition).

**Figure 2.**
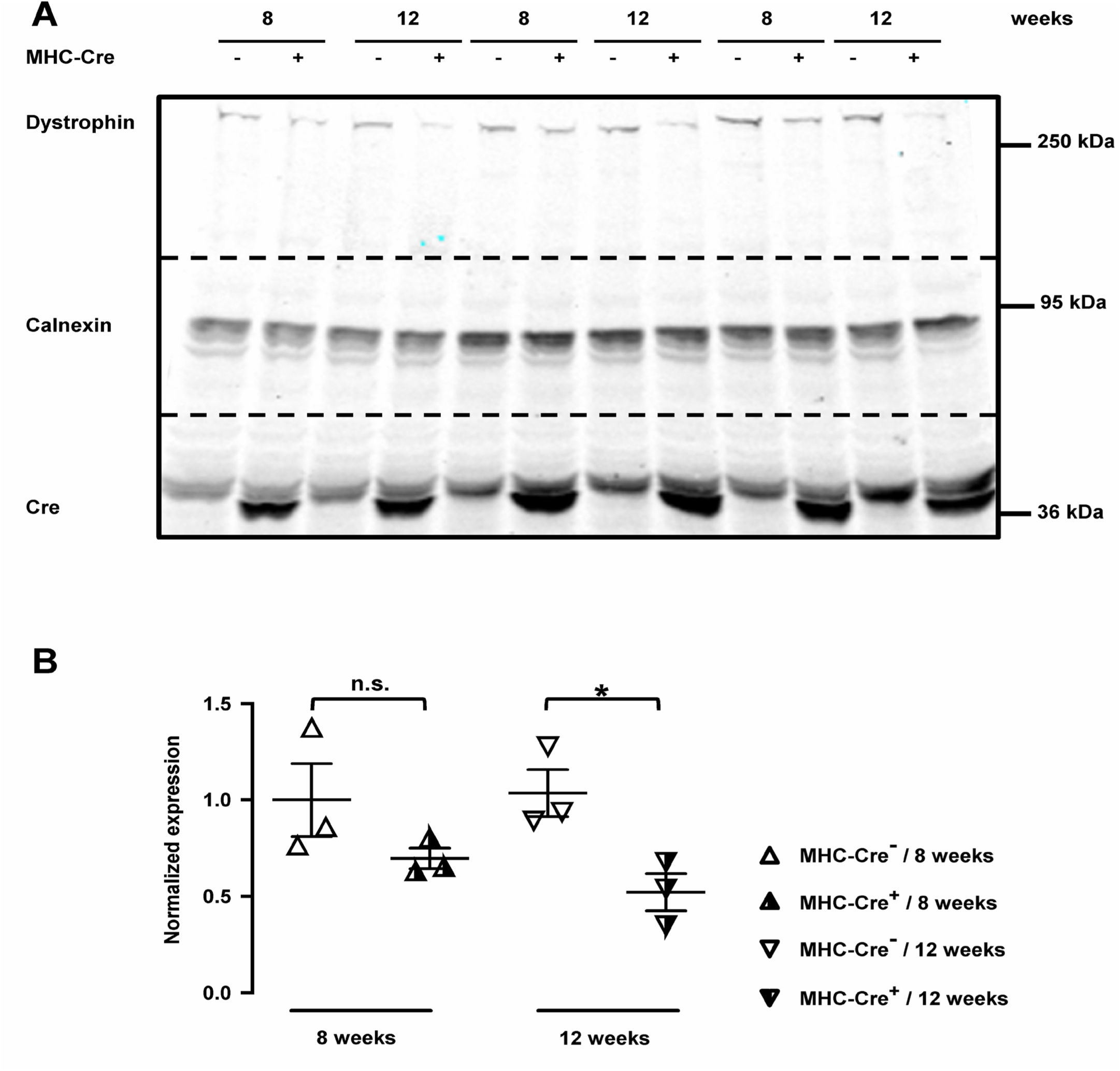
Dystrophin expression is decreased in MHC-Cre^+^ mouse hearts in an age-dependent manner. A) Western blots showing dystrophin expression in MHC-Cre^-^ and MHC-Cre^+^ mouse hearts from 8-week-old and 12-week-old mice. B) Quantification of dystrophin expression from western blots shown in A (N=3 for each condition). * p < 0.05, one-way ANOVA followed by Sidak's multiple comparisons test.

**Figure 3.**
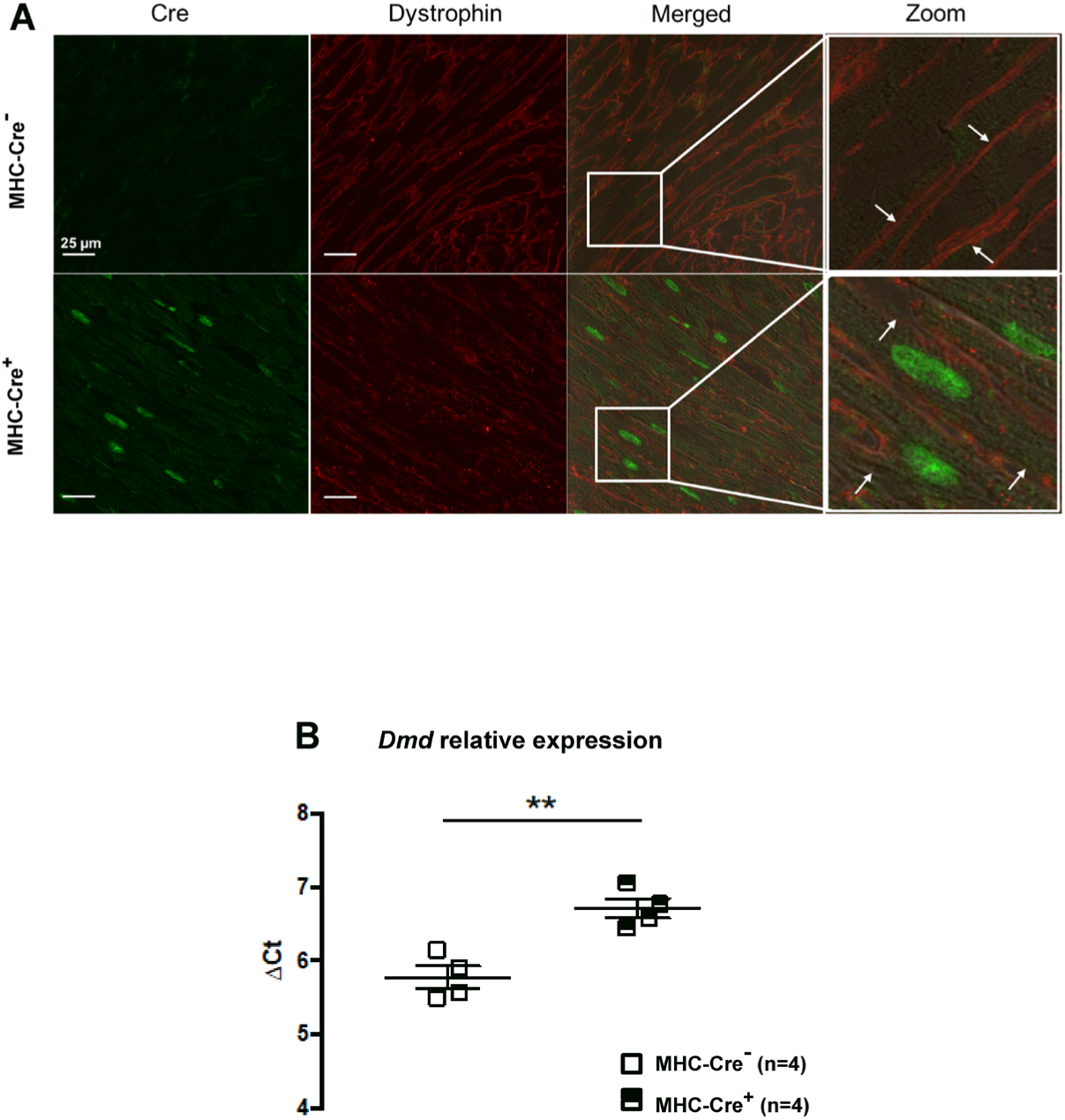
Dystrophin expression is altered in adult MHC-Cre^+^ mouse hearts at protein and mRNA level. A) Immunostainings showing decreased dystrophin signal (red) at the lateral membrane (white arrows), in cardiomyocytes expressing Cre (green) (N=3 for each condition). B) Decreased mRNA levels for dystrophin (increased ΔCT) are also observed in adult MHC-Cre^+^ mouse hearts compared with control littermates (MHC-Cre^-^) (N=4 for each condition). ** p < 0.01, t-test two tailed.

### 3.2 Dmd RNA expression is reduced in MHC-Cre hearts

To investigate at which level Cre affects dystrophin protein expression, we performed quantitative RT-PCR using RNAs extracted from >12-week-old mouse hearts. As shown in figure 3B, the relative increase of ΔCt measured with MHC-Cre^+^ hearts correspond to a ~50% reduction in *Dmd* mRNA level in MHC-Cre^+^ compared to the control mice (MHC-Cre^-^) (Fig. 3B). This result suggests that Cre decrease mRNA expression by an unknown mechanism.

### 3.3 Calcium current I_CaL_ but not Na_v_1.5 expression is altered in MHC-Cre^+^ cardiomyocytes

Considering the role of dystrophin in the function and regulation of various ion channels, we recorded calcium currents in isolated cardiomyocytes of Cre-expressing cardiomyocytes. Cardiomyocytes from 12-week old MHC-Cre^+^ mice showed a decreased current density in compared to MHC-Cre^-^ (Fig. 4A). Since dystrophin regulates sodium channel expression in cardiac myocytes, we also studied Na_v_1.5 expression in whole-heart lysates with western blots. Interestingly, Na_v_1.5 expression did not differ between MHC-Cre^+^ hearts compared to controls, neither in young nor older adult hearts (Fig. 4B).

**Figure 4.**
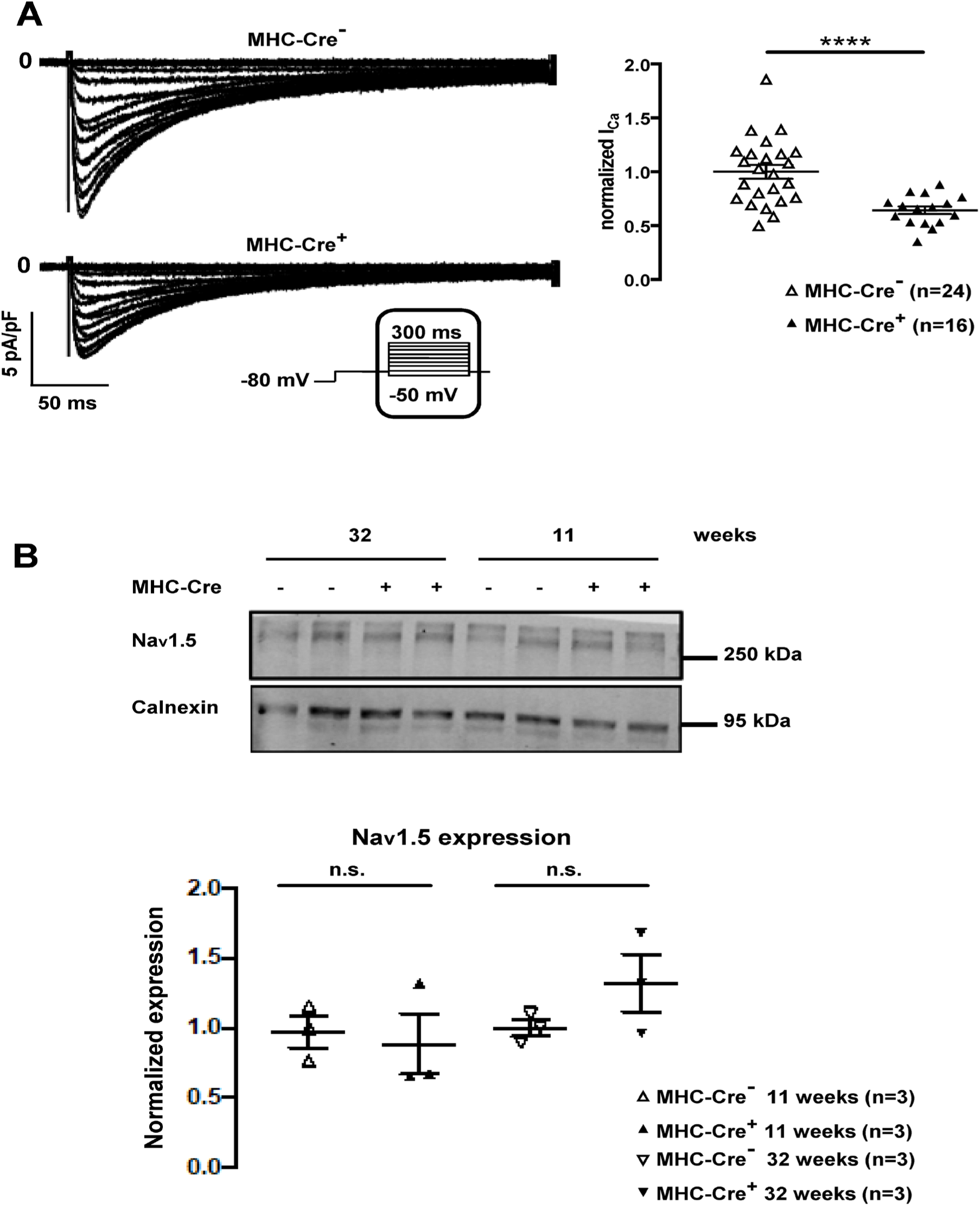
Decreased I_CaL_ and unaltered Na_v_1.5 expression in MHC-Cre^+^ mouse hearts. A) Representative traces of I_CaL_ recorded in isolated cardiomyocytes (left panel) and quantification of I_CaL_ (right panel) shown a significant down-regulation of the calcium current in MHC-Cre^+^ cardiomyocytes compared to the control (MHC-Cre^-^)(MHC-Cre^+^ N=4, n=24 and MHC-Cre^-^ N=4, n=16)). **** p < 0.0001, Welch's t-test (MHC-Cre^-^ n=24 and MHC-Cre^+^ n=16). B) Western blots showing Na_v_1.5 expression in heart lysates from young and old adult MHC-Cre^+^ and MHC-Cre^-^ mice. The bottom panel shows the quantification of Na_v_1.5 expression normalized to calnexin expression; n.s. indicates a non-statistically significant difference; one-way ANOVA followed by Sidak's multiple comparisons test (N=3 for each condition).

### 3.4 The TNT-Cre mouse line may be a better alternative model

To check whether another cardiac-specific Cre-expressing mouse line also shows a reduction in dystrophin expression, western blots were performed using heart lysates from TNT-Cre^+^ mice, a mouse line where Cre recombinase is under the control of the rat troponin T2 cardiac promoter. This strain has been a useful tool for generating early cardiomyocyte-specific mutants as Cre is mainly expressed during embryogenesis^9^. As shown in Fig. 5, no decrease in dystrophin protein expression was observed in old adult TNT-Cre^+^ mouse hearts compared to control littermates (TNT-Cre^-^) (Fig. 5A). In old adult TNT-Cre^+^ mice, moreover, neither Na_v_1.5 expression (Fig. 5B) nor calcium currents (Fig. 5C) were altered in comparison with control heart lysates and cardiomyocytes. Interestingly, Cre expression could not be observed in adult TNT-Cre hearts. Cre expression was however noticeable in heart lysates from newborn TNT-Cre^+^ mice (Suppl. Fig. 1B). In addition, when TNT-Cre^+^ mice were crossed with either *Snta1*-floxed or *Dlg1*-floxed genes, coding respectively for α1-syntrophin and the MAGUK protein SAP97, both proteins were down regulated. A similar down-regulation was observed in MHC-Cre^+^ mice in which either *Snta1* or *Dlg1* was floxed. These observations confirm that Cre functions properly in both TNT-Cre and MHC-Cre mouse lines (Suppl. Fig. 2A and 2B).

**Figure 5.**
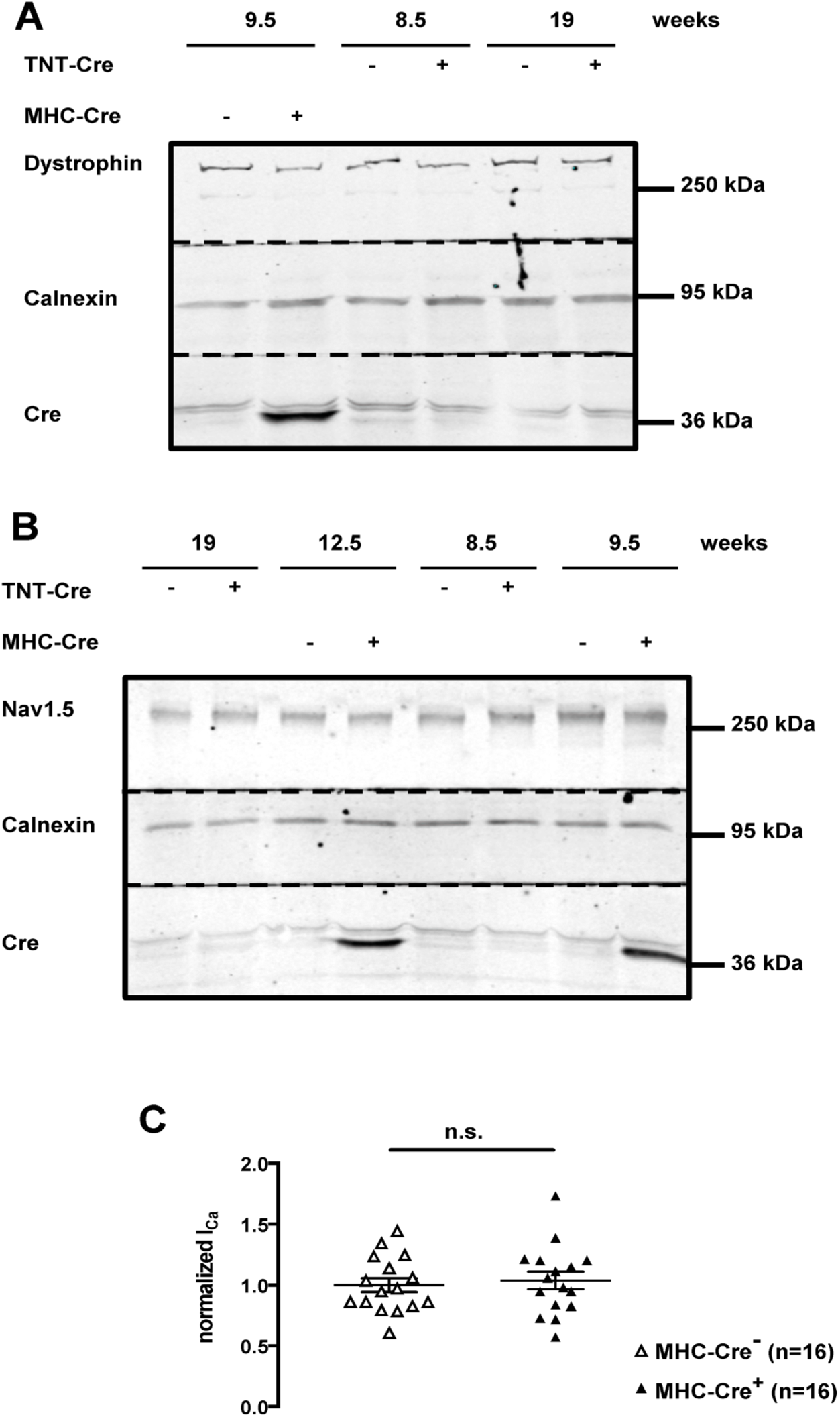
Characterization of adult TNT-Cre^+^ and (-) mouse hearts. A) Western blots showing dystrophin expression in adult MHC-Cre^+^ and MHC-Cre^-^ and TNT-Cre^+^ and TNT-Cre^-^ mouse heart lysates (N=3 for each condition). B) Western blots showing Na_v_1.5 expression in adult TNT-Cre^+^ and TNT-Cre^-^ and MHC-Cre^+^ and MHC-Cre^-^ mouse heart lysates (N=3 for each condition). C) Calcium current is not altered in isolated cardiomyocytes from TNT-Cre^+^ mouse hearts compared to TNT-Cre^-^ control hearts (MHC-Cre^-^ N=4, n=16 and MHC-Cre^+^ N=4, n=16).

## Discussion

MHC-Cre mice have been extensively used to alter the expression of various cardiac genes^10–12^. Our present work however strongly suggests that the MHC-Cre model should be carefully characterized before using it for any experiment. This recommendation stems from our following findings. Firstly, Cre expression alone reduces dystrophin protein and mRNA expression from about 12 weeks of age, while dystrophin is a crucial protein in many cardiac processes, and dystrophin reduction has been observed in human diseased cardiomyocytes^4^. Secondly, the L-type calcium current density is decreased by about 40%. Calcium current is pivotal for cardiac excitation-contraction coupling, among others, and calcium current dysregulation has been implied in many cardiac arrhythmias. Cre expression alone therefore likely induces a disease-like phenotype, which may cloud the effects of the deletion of any other specific gene a researcher may target. Indeed, Buerger *et al.* have observed dilated cardiomyopathy in MHC-Cre^+^ mice, further highlighting a potential deleterious effect of the Cre recombinase^12^. In 2015, Pugach *et al*. also reported that prolonged expression of Cre recombinase under MHC decreases cardiac function and leads to DNA damage, fibrosis, and cardiac inflammation^3^. Given that the constitutive expression of Cre in the MHC-Cre model may underlie the aforementioned problems, an αMHC-MerCreMer mouse line has been generated^13^. MerCreMer is a fusion protein containing Cre recombinase with two modified estrogen receptor ligand-binding domains at both ends. This model also allows the cardiac-specific expression of Cre recombinase driven by the MHC promoter, but provides an alternative to the constitutive expression of the Cre enzyme in the MHC-Cre^+^ model. To induce Cre expression in cardiomyocytes, MHC-MerCreMer mice have to be treated with estrogen receptor modulators such as tamoxifen. Stec *et al*. however reported that the systolic function is impaired in MHC-MerCreMer mouse strain^14^. In 2013, two groups moreover showed that the expression of Cre recombinase after induction with tamoxifen is linked to cardiac fibrosis and DNA damage leading to heart failure and death^15,16^. Recently, raloxifene, another estrogen receptor activator, has been proposed as a replacement for tamoxifen. Raloxifene induces Cre expression in a similar way, and no cardiac dysfunctions have been reported as yet^17^. However, the complexity of the administration procedure of raloxifene limits its utilization. Another proposed strategy is to regulate Cre recombinase expression with a reverse tetracycline transactivator, specifically expressed in the heart^18^. Using this approach, doxycycline added to food and drink inhibits the Cre expression^18^. Once doxycycline is removed, the Cre enzyme is produced. However, to our knowledge, the effect of this approach on cardiac function has not been investigated yet. Recently Werfel *et al*. established a promising approach for cardiac gene inactivation by transfer of the Cre recombinase gene using adeno-associated viral vectors serotype 9^19^. Such technology may allow the strong inactivation of the floxed gene specifically in cardiac tissue without adverse side effects on cardiac functions^19^.

In the present study, we unexpectedly observed that expression of the sodium channel Na_v_1.5 is not affected by Cre expression, despite the role of dystrophin in the stabilization of Na_v_1.5 at the lateral sarcolemma, which has been demonstrated in dystrophin-deficient mdx^5cv^ mice 5. One may speculate that, because dystrophin expression is lost at an earlier time point in mdx^5cv^ than in MHC-Cre^+^ mice, Na_v_ 1.5 is not yet affected in MHC-Cre^+^ mice. In addition, an increase in utrophin expression may compensate for the dystrophin down-regulation, which has already been reported in previous studies^20–22^. Importantly, in a recent study, the presence of more than 600 loxP like sites in the genome of C57Bl/6 mice has been reported^3^. Based on this observation, one may propose that many genes of the MHC-Cre^+^ mouse strain can be modified, leading to various unexpected effects. As of yet, no comprehensive proteomic analysis has been performed on MHC-Cre^+^ model, we may speculate that other genes and proteins are dysregulated in this model, which could compensate for the down-expression of Na_v_1.5 channels. Surprisingly, the expression of Cre recombinase under the rat troponin T2 cardiac promoter affected neither dystrophin expression nor the calcium current. However, a critical limitation for the interpretation of this finding is that the MHC-Cre^+^ and TNT-Cre^+^ mice differed in their mixed genetic background levels. As already reported, the background of the mice is important for the onset of cardiac dysfunctions due to Cre recombinase expression^23^. Mice on mixed backgrounds seem to be more resistant to developing cardiac dysfunction in response to Cre expression^23^. Consequently, it is important to repeat these experiments with MHC-Cre^+^ and TNT-Cre^+^ mice with mice on a pure background.

In conclusion, the findings of the present study are a clear warning to all researchers who use engineered mouse lines, especially MHC-Cre. We strongly recommend keeping our results in mind when designing experimental procedures involving MHC-Cre mice, and probably other Cre-expressing mouse models. Experimenters should carefully choose the proper controls, and consider the age, and likely the sex of the animals.

## Acknowledgments

We are very grateful to Stephan Sonntag and Doron Shmerling from Polygene, for the generation of the *Snta1*^flox^, *Dmd*^flox^, and *Dlg1*^flox^ mouse strains, and to Sarah Vermij for editing and proofreading.

## Data Availability

“All data underlying the results are available as part of the article and no additional source data are required.”

## SUPPLEMENTARY MATERIAL

**Supplemental Figure 1.**
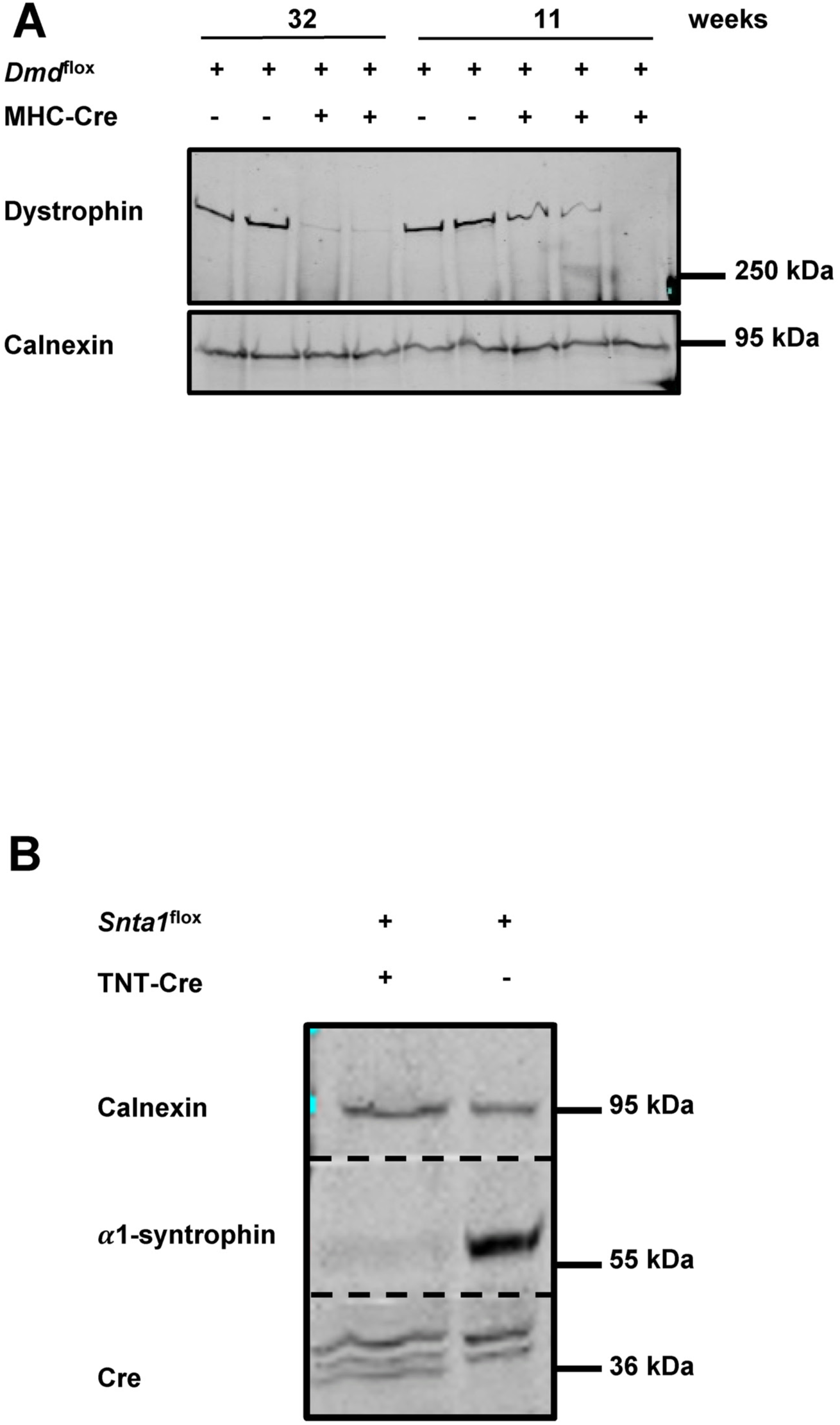
A) Western blots showing the decreased dystrophin expression in adult hearts of *Dmd^flox^* mice with Cre recombinase (MHC-Cre^+^) or without (MHC-Cre^-^) (N= 2 to 3 for each condition). B) Western blots showing Cre recombinase and α1-syntrophin expression in neonatal hearts from *Snta1*^flox^ with Cre under control of the TNT promoter (TNT-Cre^+^) or without (TNT-Cre^-^) (N=2 for each condition).

**Supplemental Figure 2.**
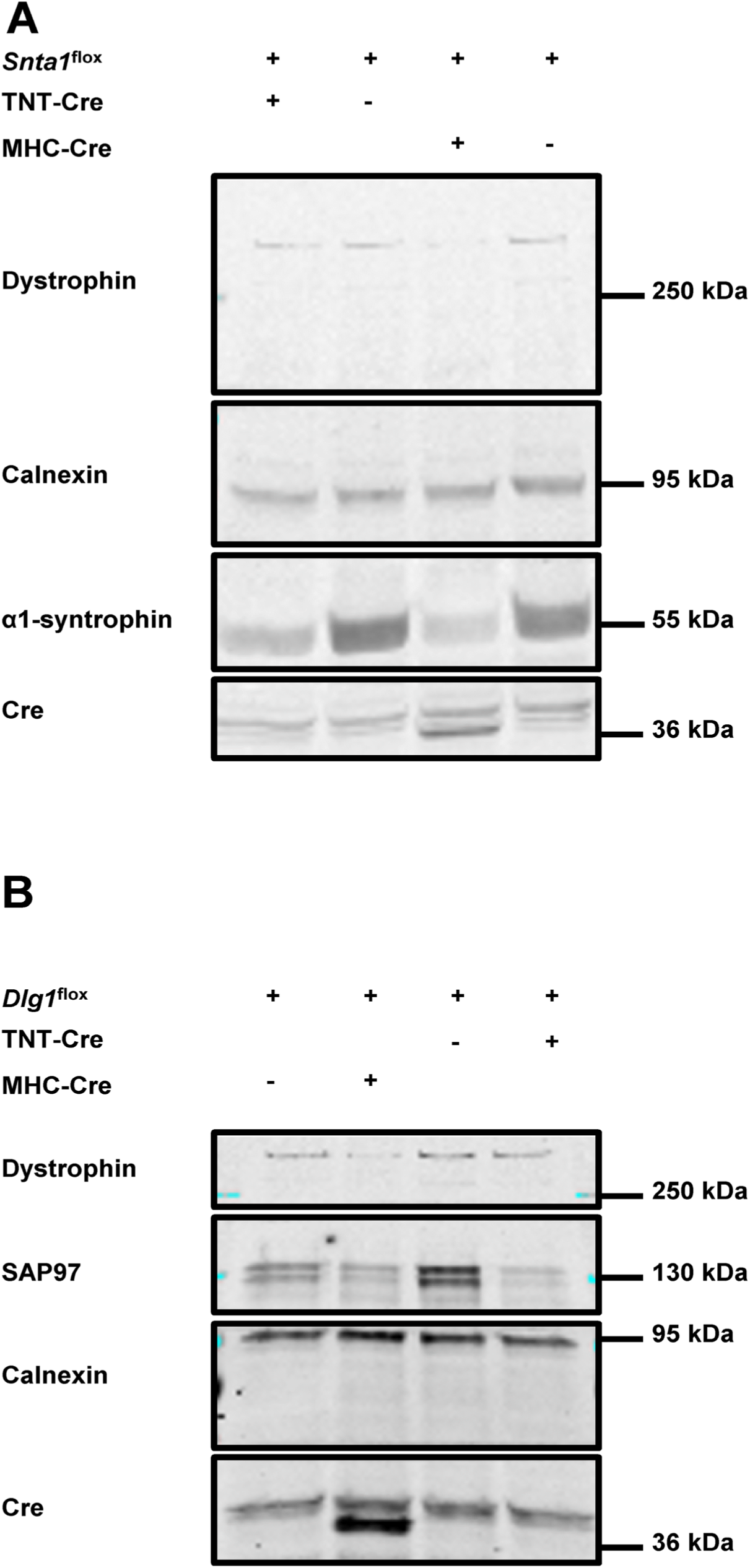
A) Western blots showing a reduction of the expression of syntrophin-α1 in adult mouse hearts using *Snta1^flox^* mice and TNT-Cre^+^ or MHC-Cre^+^ mouse strains (N=3 for each condition). B) Western blots showing the decrease of expression of SAP97 in adult mouse hearts using *Dlg1*^flox^ mice and TNT-Cre^+^ or MHC-Cre^+^ mouse strains (N=3 for each condition).

